# Age-related losses in cardiac autonomic activity during a daytime nap

**DOI:** 10.1101/2020.06.26.168278

**Authors:** Pin-Chun Chen, Negin Sattari, Lauren N. Whitehurst, Sara C. Mednick

## Abstract

In healthy, young individuals, a reduction in cardiovascular output and a shift from sympathetic to parasympathetic (vagal) dominance is observed from wake into stages of nocturnal and daytime sleep. This cardiac autonomic profile, measured by heart rate variability (HRV), has been associated with significant benefits for cardiovascular health. Aging is associated with decreased nighttime sleep quality and lower parasympathetic activity during both sleep and resting. However, it is not known whether age-related dampening of HRV extends to daytime sleep, diminishing the cardiovascular benefits of naps in the elderly. Here, we investigated this question by comparing the autonomic activity profile between young and older healthy adults during a daytime nap and a similar period of wakefulness (quiet wake; QW). For each condition, from the electrocardiogram (ECG), we obtained beat-to-beat HRV intervals (RR), root mean square of successive differences between adjacent heart-beat-intervals (RMSSD), high frequency (HF), low frequency (LF) power and total power (TP), HF normalized units (HFnu), and the LF/HF ratio. As previously reported, young subjects showed a parasympathetic dominance during NREM, compared with REM, pre-nap rest, and WASO. On the other hand, older, compared to younger, adults showed significantly lower vagally-mediated HRV (measured by RMSSD, HF, HFnu) during NREM. Interestingly, however, no age-related differences were detected during pre-nap rest or QW. Altogether, our findings suggest a sleep-specific reduction in parasympathetic modulation that is unique to NREM sleep in older adults.

**Impact Statement:** Sleep is naturally modulated by the autonomic nervous system (ANS), with greater dominance of parasympathetic over sympathetic activity during non-rapid-eye-movement (NREM) sleep. As such, sleep has been termed a “cardiovascular holiday” and has been associated with positive health outcomes. Aging, however, is linked to decreases in cardiac autonomic activity and sympathovagal imbalance. While the impact of aging on ANS activity during nocturnal sleep has received some attention, the cardiac profiles during a daytime nap, to our knowledge, have not yet been studied under the context of aging. Herein, young adults demonstrated increased parasympathetic activity during deep sleep. Older adults, however, showed less parasympathetic modulation during NREM sleep, suggesting loss of the cardiovascular holiday. Importantly, no age-related declines in parasympathetic activity were detected during wake, suggesting a sleep-specific reduction in parasympathetic modulation that is unique to NREM sleep in older adults.

## Introduction

The autonomic nervous system (ANS) plays a major role in cardiovascular adaptations to environmental changes and is thus critical for global allostasis (McEwen, 2002), or the ability of organisms to maintain stability, adaptability, and health in response to actual or perceived environmental and psychological demands. The ANS is generally conceived to have two major branches—the sympathetic system, associated with energy mobilization, and the parasympathetic system, associated with vegetative and restorative functions. In healthy adults, the activity of these branches is in dynamic balance and is dependent on activity and rest cycles. For example, there is a well-documented circadian rhythm to the ANS such that sympathetic activity is higher during daytime hours and parasympathetic activity increases at night (Jarczok et al., 2019). Along with these circadian shifts, sleep also directly influences the ANS, and vice versa, with restorative, parasympathetic processes dominating during nighttime sleep (Trinder et al., 2012). In younger adults, daytime naps present with similar cardiac autonomic profiles as nighttime sleep (Cellini et al., 2016; Whitehurst et al., 2018), suggesting that daytime sleep too, serves a restorative function. These circadian- and sleep-dependent shifts in the ANS are critical to the maintenance of autonomic balance between parasympathetic and sympathetic branches (Boudreaux, 2013) and are beneficial for health and cognition (Thayer et al. 2009). As we age, ANS functions can become more rigid, with decreased β-adrenoreceptor response, decreases in baroreflex activity, diminished heart rate responses to acetylcholine, and decreases in cardiac parasympathetic activity (Chadda et al., 2018). Studies show that these age-related ANS changes may contribute to autonomic imbalance, in which the sympathetic system is hyperactive and/or the parasympathetic system is hypoactive. Autonomic imbalance is associated with various pathological conditions, such as cardiovascular morbidity and mortality, diabetes, and dementia (Thayer et al., 2010; Thayer & Lane, 2007). It is not yet known whether cardiac autonomic profiles during daytime naps differ in both older and younger adults, which was the goal of the present investigation. Such information would contribute to a comprehensive understanding of sleep’s role in ANS activity across the life span.

One way to determine ANS activity is through heart rate variability (HRV) measurements that reflect global ANS changes, and specifically, parasympathetic activity. Parasympathetic/vagal activity is implicated in two measurements extracted from the electrocardiogram (ECG). First, is the root mean square of successive differences between adjacent heart beat-to-beat intervals (RMSSD) where higher intervals between heart beats represents greater parasympathetic/vagal innervation (Task Force, 1996). The second is high frequency power (HF; 0.15–0.40 Hz), which is indicative of vagally mediated respiration (Task Force, 1996). Decreased parasympathetic activity is associated with excessive inflammatory functioning (Pavlov & Tracey, 2012), and heightened risk for chronic disease (see Kemp & Quintana, 2013 for a review), including cardiovascular disease, which is the leading cause of death and disability in the United States (CDC, 2017). Beyond physical health, HRV has also been linked to greater self-regulation of cognitive, emotional, and social domains (see Shaffer et al., 2014 for a review). For example, Thayer and colleagues proposed the Neurovisceral Integration Model and argued that HRV reflects the activity of an integrative neural network that flexibly regulates physiological, cognitive, and emotional responses (Smith et al., 2017; Thayer & Lane, 2009). Specifically, individuals with higher HRV during wake (measured during or immediately prior to cognitive testing) have been shown to perform better on a wide range of cognitive tasks that engage the prefrontal cortex, including working memory (n-back task: Hansen et al., 2003; operation-span task: Mosley et al., 2018), cognitive inhibition (Stroop task: Hansen et al., 2004), and emotion regulation (Williams et al., 2015). Considering HRV may be a critical predictor of psychological and physiological health, HRV can be considered an access point for intervention as sleep is a reoccurring condition where ANS activity is naturally modulated, and importantly, good sleep has been tied to better physical and cognitive health outcomes (see Whitehurst et al., 2020 for a review).

Sleep and the ANS influence each other in a bidirectional fashion. Changes in the ANS modulate sleep onset and the transition between the different sleep stages, and each sleep stage is associated with a distinct autonomic profile (see Trinder et al., 2012 for a review). As the brain shifts from wake to sleep, the body undergoes marked changes, including heart rate deceleration and relative increases in parasympathetic activity during non-rapid eye movement (NREM) sleep (Trinder et al., 2001). Young adults show variations in HRV according to sleep stages, switching from vagally dominated NREM (N2 and N3) sleep to sympathetic-parasympathetic balanced rapid eye movement (REM) sleep (Brandenberger et al, 2001). ANS activity during nocturnal sleep has been considered as a “cardiovascular holiday” (see Tobaldini et al., 2013 for a review) due to its characteristic drop in blood pressure and parasympathetic dominance during NREM sleep compared with wake. However, whether such cardiovascular benefits are solely sleep-dependent has received some challenges.

Circadian rhythms influence both sleep (Borb & Achermann, 1999) and cardiovascular activity (Guo & Stein, 2003), and may contribute to promoting parasympathetic activation. As such, daytime sleep may confer less restorative benefit (i.e., parasympathetic tone) than nighttime sleep. However, few studies have investigated cardiac profiles during daytime naps. Two recent studies report that, in healthy young subjects, vagal activity is modulated by daytime sleep (Cellini et al., 2016), and that similar HRV profiles exist between naps and nighttime sleep (Whitehurst et al., 2018). These studies support the notion that naps, like nighttime sleep, may serve as a mini-cardiovascular break. Further, and aligned with the Neurovisceral Integration Model, studies have shown that parasympathetic activity during a daytime nap facilitates improvement in long-term memory (Whitehurst et al., 2016) and working memory (Chen et al., 2020). Furthermore, these previous investigations did not directly address whether cardiovascular benefits during a nap are sleep-specific or whether benefits can also be achieved from a comparable duration of quiet wake during the day. Specifically, these studies typically compared the profile across the nap sleep stages to a five-minute pre-sleep rest period, reporting increased parasympathetic activity during NREM sleep compared with pre-sleep rest. Yet, parasympathetic dominance can be induced during periods of quiet wake not associated with sleep. For example, mindfulness has been shown to enhance parasympathetic influences on the heart rate (increased HF and decreased LF; Heckenberg et al., 2018; Mankus et al., 2013; Krygier et al., 2013). In addition, listening to classical music for 20 minutes also increases parasympathetic tone (increased HF and RMSSD; Lin et al., 2013). Complicating matters, methods for measuring awake resting HRV varies between studies. Sleep studies measure HRV during five minutes of pre-sleep rest while subjects lay on a bed preparing for sleep (Cellini et al., 2016), while waking studies measure it in subjects who are seated instead of lying down, but not preparing for sleep (Mattarozzi et al., 2019). One goal of the current study is to determine if QW and pre-sleep rest confer the same cardiac profile, information that will greatly inform our understanding of HRV activity across waking states and methodologies.

Aging is linked to a decline in cardiovascular control (Colosimo et al., 1997) and substantial research shows that aging is associated with a decrease in HRV and sympathovagal imbalance (O’Brien et al., 1986; Russoniello, 2013; Santillo et al., 2012). The age-related reduction in resting HRV has been observed in both cross-sectional comparisons and in a longitudinal study (Sinnreich et al., 1998). In particular, a study found two significant declines in waking ANS activity across aging, one around adolescence/early adulthood and the second at about 60 years old with continual decline to 80 years old (Zhang, 2007). But, studies on age-related HRV changes during sleep remain scarce. The two studies that do exist report a loss of parasympathetic activity (measured by HF HRV and SDNN; standard deviation of normal to normal RR intervals) during nocturnal NREM sleep in the elderly compared with young subjects, with no significant sleep-stage dependent variations (Brandenberger et al., 2003; Crasset et al., 2001). These results imply that decreased HRV in older adults may be especially prominent during NREM sleep stages, compared to waking. The age-related cardiac profiles during a daytime nap, to our knowledge, have not yet been studied. Given that cardiovascular disease is the leading cause of death in the United States (Benjamin et al., 2017), the lack of research examining the relationship between napping and cardiovascular modulation is surprising considering that 90% of adults 60+yrs in America nap at least once a week (and 30% nap > twice a week), and that the frequency of napping has consistently been reported to increase with advancing age (Ohayon & Vecchierini, 2002; Ohayon & Zulley, 1999). Additionally, the relationship between napping and cardiovascular risk is quite controversial, with epidemiological studies reporting that frequent napping is associated with both increased (Jung, Song, Ancoli-Israel, & Barrett-Connor, 2013; Leng et al., 2014) and decreased (Campos & Siles, 2000) risk for coronary heart disease or cardiovascular mortality. Moreover, both positive (Blackwell et al., 2006; Cross et al., 2015) and negative (Campbell et al., 2005; Keage et al., 2012) associations have emerged between napping and increased risk of cognitive decline.

Given the lack of studies investigating HRV across daytime sleep in older populations and the pressing need for understanding the role of napping on physiological and psychological health, the current study aimed to assess cardiac autonomic activity across sleep stages during an afternoon nap in healthy young and older adults. First, we utilize a between-subject design to compare HRV profiles between a standard HRV QW condition and pre-nap rest condition. Next, we utilize a within-subject design to compare HRV profiles during pre-nap rest and sleep stages. We utilized a nap period of ~ 90 minutes that was strategically positioned in the mid-afternoon (1:30 pm) to promote adequate proportions of N2, N3 and REM sleep (Mednick et al., 2003; Mednick et al., 2013). Our a priori hypotheses were that: (1) QW will show similar parasympathetic activity compared to pre-nap rest in both age groups; (2) young adults will show parasympathetic dominance during NREM sleep compared to pre-nap rest; (3) older adults will show no significant difference in parasympathetic modulation during NREM sleep compared to pre-nap rest; (4) older adults will show decreased parasympathetic modulation during NREM sleep and QW, compared to the younger subjects.

## Methods

### Participants

60 (NAP)+46 (QW) healthy young adults (18-35yo, male=60, Mean =20.67, SD = 3.07) and 46 (NAP)+38 (QW) older adults (60-85yo, male=42, Mean = 69.15, SD = 5.93) with no personal history of neurological, psychological, or other chronic illness provided informed consent, which was approved by the University of California, Riverside Human Research Review Board. Demographics and prior self-reported sleep habits were reported in Table 1. Participants included in the study had a regular sleep wake schedule (reporting a habitual time in bed of about 7–9 h for young adults and 6-8 h for older adults per night; National Sleep Foundation, 2015).

**Table 1.**
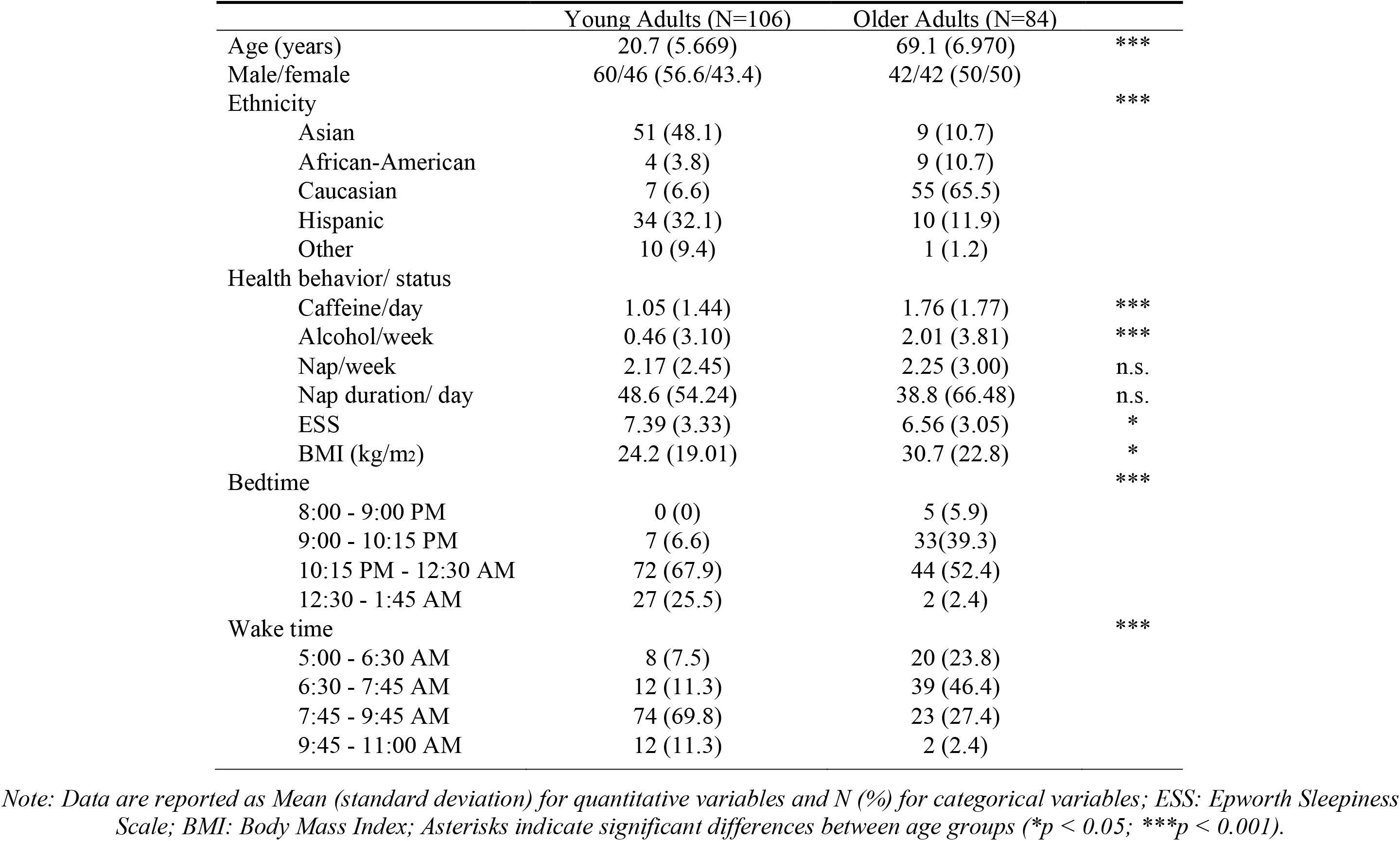
Descriptive statistics for Demographics

|  | Young Adults (N=106) | Older Adults (N=84) |  |
| --- | --- | --- | --- |
| Age (years) | 20.7 (5.669) | 69.1 (6.970) | *** |
| Male/female | 60/46 (56.6/43.4) | 42/42 (50/50) |  |
| Ethnicity |  |  | *** |
| Asian | 51 (48.1) | 9 (10.7) |  |
| African-American | 4 (3.8) | 9 (10.7) |  |
| Caucasian | 7 (6.6) | 55 (65.5) |  |
| Hispanic | 34 (32.1) | 10 (11.9) |  |
| Other | 10 (9.4) | 1 (1.2) |  |
| Health behavior/ status |  |  |  |
| Caffeine/day | 1.05 (1.44) | 1.76 (1.77) | *** |
| Alcohol/week | 0.46 (3.10) | 2.01 (3.81) | *** |
| Nap/week | 2.17 (2.45) | 2.25 (3.00) | n.s. |
| Nap duration/ day | 48.6 (54.24) | 38.8 (66.48) | n.s. |
| ESS | 7.39 (3.33) | 6.56 (3.05) | * |
| BMI (kg/m<sup>2</sup>) | 24.2 (19.01) | 30.7 (22.8) | * |
| Bedtime |  |  | *** |
| 8:00 - 9:00 PM | 0 (0) | 5 (5.9) |  |
| 9:00 - 10:15 PM | 7 (6.6) | 33(39.3) |  |
| 10:15 PM - 12:30 AM | 72 (67.9) | 44 (52.4) |  |
| 12:30 - 1:45 AM | 27 (25.5) | 2 (2.4) |  |
| Wake time |  |  | *** |
| 5:00 - 6:30 AM | 8 (7.5) | 20 (23.8) |  |
| 6:30 - 7:45 AM | 12 (11.3) | 39 (46.4) |  |
| 7:45 - 9:45 AM | 74 (69.8) | 23 (27.4) |  |
| 9:45 - 11:00 AM | 12 (11.3) | 2 (2.4) |  |

*Note: Data are reported as Mean (standard deviation) for quantitative variables and N (%) for categorical variables; ESS: Epworth Sleepiness Scale; BMI: Body Mass Index; Asterisks indicate significant differences between age groups (\* $p < 0.05$ ; \*\*\* $p < 0.001$ ).*

**Table 2.** Descriptive statistics for nap

|  | Young Adults | Older Adults |  |
| --- | --- | --- | --- |
| TIB (min) | 78.069 (3.1) | 80.763 (3.36) |  |
| TST (min) | 61.500 (2.94) | 46.987 (3.92) | ** |
| SOL (min) | 7.664 (0.99) | 9.118 (1.27) |  |
| N1 (min) | 4.586 (0.47) | 7.934 (1.07) | ** |
| N2 (min) | 31.448 (1.94) | 28.855 (2.86) |  |
| N3 (min) | 19.224 (1.96) | 7.474 (1.58) | *** |
| REM (min) | 6.241 (1.09) | 2.72 (0.98) |  |
| WASO (min) | 6.560 (0.88) | 19.01 (2.12) | *** |
| SE (%) | 77.588 (2.21) | 56.52 (3.47) | *** |

*Note: Data are reported as Mean (standard error); TIB = Total Time in Bed; TST = Total Sleep Time; N3 = Slow Wave Sleep; REM = Rapid Eye Movement sleep; WASO = Wake After Sleep Onset (calculated as the minutes of wake after first epoch of sleep); SOL = Sleep Latency (calculated as the time to first epoch of sleep); SE = Sleep Efficiency (calculated as $100 \times TST/TIB$ ). All stats are represented in minutes besides SE which is in percentage. Asterisks indicate significant differences between age groups (\*\* $p < 0.01$ ; \*\*\* $p < 0.001$ ).*

The personal health histories were measured twice. First, during a pre-screening questionnaire in which a general health condition report was collected over the online survey and the eligibility was determined. Second, eligible subjects were invited to participate in an orientation in which they were given details about the study and interviewed by a trained graduate student. Participants who met any of the following exclusion criteria were excluded from the study: a) extreme morning or evening-type tendencies (measured with the Morningness Eveningness Questionnaire; Horne & Östberg, 1976); b) excessive daytime sleepiness (reported by Epworth Sleepiness Scale; Johns, 1991; subjects' rating > 13 were excluded); c) a sleep disorder (assessed by questionnaires); d) any personal or immediate family (i.e., first degree relative) history of diagnosed significant psychopathology; e) personal history of head injury with loss of consciousness greater than 2 minutes or seizures; f) history of substance dependence; g) current use of any psychotropic medications; h) any cardiac, respiratory or other medical condition which may affect cerebral metabolism; i) non-correctable vision and auditory impairments.

Participants who did not met any of the exclusion criteria above and also met all the following inclusion criteria were enrolled in the study: a) aged 18-39/ 60-85 years old; b) healthy, non-smoking adult without major medical problems; c) completed at least 12 years of education; d) a regular sleep-wake schedule, defined as obtaining 7–9 h (young adults) or 6-8 h (older adults) of sleep per night, with a habitual bedtime between 9pm and 2am (young adults) or 8pm and 1am (older adults) and a habitual wake time between 6am and 10am (young adults) or 5am and 9am (older adults). Enrolled participants were asked to maintain their schedule for one week prior to their visit, which was monitored with sleep diaries. In addition, participants were asked to wear an actigraph (Actiwatch Spectrum, Respironics) for one night prior to their visit. Subjects were rescheduled if they reported poor sleep quality in their sleep diary, such as having more than 2 nights of less than 6 h of sleep or more than 9 h of sleep during the week prior to their visit, or if subjects’ actigraphy data showed less than 6 h of sleep or more than 9 h of sleep the night before the experimental visit. Rescheduled subjects were given another week to fill out a new sleep diary and maintain a regular sleep wake schedule prior to their visit.

During the orientation, participants were screened for cognitive impairment using Digit Span Backwards, which contains a multi-variate length string of digits; subjects were asked to repeat the string of digits backwards after it was read to them. In addition, Older participants were screened for dementia using the Telephone Screening for Dementia questionnaires (referred to as TELE) (Gatz et al., 2002), which consists of variety of questions to identify dementia-like symptoms. For older subjects, the STOP-BANG questionnaire (Chung et al., 2008) was used to screen for obstructive sleep apnea and those with a medium to high risk (answering yes to three or more items) were excluded. Additionally, participants were instructed to abstain from caffeine and alcohol starting at noon the day prior to the study (detected on the sleep diary). Participants who consumed alcoholic beverages > 10 cans a week or caffeinated products > 3 cups a day were excluded prior to the study (reported in Table 1).

### Procedures

On the study day, subjects were randomly assigned to either a nap condition (NAP) or a quiet wake condition (QW). For the NAP group, subjects were provided a 2-hour nap opportunity and were allowed to sleep up to 90 min, monitored with polysomnography (PSG), including electroencephalography (EEG), electrocardiogram (ECG), electromyogram (EMG), and electrooculogram (EOG), in the Sleep and Cognition (SaC) lab at 1:30 pm (12:30-1:00pm for older subjects because of an assumption that their circadian rhythm and sleep schedule may be shifted forward; Monk, 2005). Before sleep, we recorded 5-min resting HRV while subjects lay awake, in a supine position with head and body still. Sleep was monitored online by a trained sleep technician. In the QW conditions, subjects watched a 50-min nature documentary in a dark room lying on the bed during which they were monitored by lab staff to make sure they did not fall sleep. Participants received monetary compensation for participating in the study.

### Sleep Recording and Scoring

Electroencephalographic (EEG) data were acquired using a 32-channel cap (EASYCAP GmbH) with Ag/AgCI electrodes placed according to the international 10-20 System (Jasper, 1958). Electrodes included 24 scalp, two electrocardiography (ECG), two electromyography (EMG), two electrooculography (EOG), 1 ground, and 1 on-line common reference channel. The EEG was recorded with a 1000 Hz sampling rate and was re-referenced to the contralateral mastoid (A1 & A2) post-recording. Only eight scalp electrodes (F3, F4, C3, C4, P3, P4, O1, O2), the EMG and EOG were used in the scoring of the nighttime sleep data. Each signal was amplified, band-pass filtered (EEG and EOG: 0.3–35 Hz), and digitized at 256 Hz. Next, data were pre-processed using BrainVision Analyzer 2.0 (BrainProducts, Munich Germany) and all epochs with artifacts and arousals were identified by visual inspection and rejected. EEG recordings were visually scored by two trained sleep technicians. Prior to scoring these data, technicians were required to reach 85% reliability with each other and one other rater in the lab on two independent data sets. Raw data were visually scored in 30-sec epochs into Wake, Stage 1 Sleep (N1), Stage 2 Sleep (N2), Slow Wave Sleep (N3) and rapid eye movement sleep (REM) according to the American Academy of Sleep Medicine (AASM) rules for sleep staging using HUME, a custom MATLAB toolbox. Minutes in each sleep stage were calculated. Sleep latency (SOL) was calculated as the number of minutes from lights out until the initial epoch of sleep, N2, N3 and REM. Additionally, wake after sleep onset (WASO) was calculated as total minutes awake after the initial epoch of sleep and sleep efficiency (SE) was computed as total time spent asleep after lights out divided by the total time spent in bed x 100.

### Heart Rate Variability

Electrocardiogram (ECG) data were acquired at a 1000-Hz sampling rate using a modified Lead II Einthoven configuration. We analyzed HRV of the R-waves series across the whole sleep/wake period using Kubios HRV Analysis Software 2.2 (Biosignal Analysis and Medical Imaging Group, University of Kuopio, Finland), according to the Task Force guidelines (Electrophysiology Task Force of the European Society of Cardiology the North American Society of Pacing, 1996). RR peaks were automatically detected by the Kubios software and visually examined by trained technicians. Incorrectly detected R-peaks were manually edited. Missing beats were corrected via cubic spline interpolation. Inter-beat intervals were computed, and a third-order polynomial filter was applied on the time series in order to remove trend components. Artifacts were removed using the automatic medium filter provided by the Kubios software.

The HRV analysis of the RR series was performed by using a Matlab-based algorithm (see Whitehurst et al., 2018). An autoregressive model (Model order set at 16; Boardman et al., 2002) was employed to calculate the absolute spectral power (ms^2^) in the LF HRV (0.04–0.15 Hz; ms2x) and the HF HRV (0.15–0.40 Hz; ms^2^; an index of vagal tone) frequency bands, as well as total power (TP; ms^2^; reflecting total HRV), and HF peak frequency (HFpf; Hz; reflecting respiratory rate). From these variables, we derived the HF normalized units (HF_nu_ = HF[ms^2^]/HF[ms^2^]+LF[ms^2^]) and the LF/HF ratio (LF[ms^2^]/HF[ms^2^]), an index often considered to reflect the sympathovagal balance (i.e., the balance between the two branches of the ANS), but whose meaning has been recently put into question (Billman, 2013; Reyes del Paso et al., 2013). The LF, HF, and TP measures had skewed distributions and as such were transformed by taking the natural logarithm, as suggested by Laborde et al., 2017. Since the LF normalized units are mathematically reciprocal to HF_nu_ (i.e. LF_nu_ =1− HF_nu_), to avoid redundancy, we computed only the HF_nu_ index, an index often thought to reflect vagal modulation (Burr, 2007). Due to controversies about the physiological mechanisms that contribute to changes in LF activity, LF, LF/HF ratio and HFnu are difficult to make for these parameters, but we report them for descriptive purposes.

In addition to the frequency domain parameters, RMSSD (ms; root mean square of successive differences) was calculated as a measure of vagally-mediated HRV in the time-domain. Similar to the frequency adjustments, to adjust for skewed distributions in the RMSSD, we report the natural logarithm. Additionally, we included the RR (ms; time interval between consecutive R-peaks, reflecting frequency of myocardial contraction) as an index of cardiac autonomic control in our analyses (Pinna et al., 2007).

For time-domain and frequency-domain HRV measures during different sleep stages, consecutive artifact-free 3-min windows of undisturbed sleep were selected across the whole nap using the following rules: (a) the 1.5-min preceding, and (b) the entire 3-min epoch selected must be free from stage transitions, arousal, or movements. Windows were identified and averaged within N2 sleep, slow-wave sleep (N3), REM sleep, and wake-after-sleep-onset (WASO). We also analyzed 5 min of pre-nap wakefulness (Rest). Epochs of N1 were not analyzed. 3-min windows were chosen given that naps have more fragmented sleep due to increased stage transitions, especially in the elderly subjects. In fact, even 3 minutes of undisturbed sleep in each sleep stage is not as common in daytime naps in the elderly population. Consecutive undisturbed 5-minute windows used by previous nocturnal sleep studies (Ako et al., 2003; Brandenberger et al., 2003; de Zambotti et al., 2014) would be implausible for analyzing HRV during naps. In addition, we analyzed artifact-free 3-min windows of 50 min QW during which the subjects watched a 50-min nature documentary in a dark room lying on the bed.

### Data Reduction

As mentioned in the above section, not everyone experienced all sleep stages with at least a stable 3-min epoch. Therefore, in the NAP group, we included 60 young and 46 older adults in analyses for Rest, 53 young and 37 older adults in analyses for N2, 42 young and 20 older adults in analyses for N3, 21 young and 6 older adults in analyses for REM sleep, as well as 43 young and 31 older adults in analyses for WASO. As we only have 6 older adults who had at least 3-min of stable REM sleep, REM data in the elderly were excluded from the analyses. In the QW group, we included 46 young and 38 older adults in the analyses. Subjects with data points greater or less than mean ± 3*standard deviation were considered as outliers and were excluded from the analyses. We excluded one older subject in the NAP group due to this criterion. Descriptive statistics for HRV variables are shown in Table 3.1 and 3.2 for the NAP and QW groups, respectively.

**Table 3.1.** Summary of HRV Parameters Across Sleep Stages

| Stage | Rest |  | WASO |  | N2 |  | N3 |  | REM |
| --- | --- | --- | --- | --- | --- | --- | --- | --- | --- |
| Age Group (N) | Young (60) | Older (46) | Young (43) | Older (31) | Young (53) | Older (37) | Young (42) | Older (20) | Young (21) |
| Epochs | 1.47 (0.366) | 1.91 (0.475) | 3.49 (0.454) | 5.41 (0.592) | 6.11 (0.389) | 4.97 (0.524) | 5.98 (0.437) | 3.75 (0.713) | 3.24 (0.618) |
| RR (ms) | 945 (19.2) | 943 (21.5) | 918 (20.2) | 901 (22.6) | 987 (19.5) | 930 (22.0) | 984 (20.0) | 928 (23.8) | 905 (22.4) |
| pNN50 (%) | 40.8 (3.02) | 13.1 (3.16) | 37.4 (3.24) | 10.8 (3.40) | 44.7 (3.08) | 16.2 (3.27) | 42.8 (3.20) | 13.3 (3.66) | 34.4 (3.70) |
| RMSSD (ln) | 4.22 (0.0805) | 4.11 (0.118) | 4.20 (0.0863) | 3.93 (0.131) | 4.28 (0.0821) | 3.85 (0.124) | 4.19 (0.0853) | 3.70 (0.146) | 4.17 (0.0985) |
| HF (ln ms<sup>2</sup>) | 6.73 (0.151) | 6.30 (0.225) | 6.74 (0.163) | 5.99 (0.253) | 6.93 (0.154) | 5.85 (0.237) | 6.73 (0.189) | 5.70 (0.283) | 6.73 (0.151) |
| LF (ln ms<sup>2</sup>) | 6.62 (0.132) | 6.22 (0.206) | 6.77 (0.155) | 5.99 (0.242) | 6.51 (0.139) | 5.76 (0.222) | 5.69 (0.151) | 5.16 (0.281) | 6.71 (0.200) |
| TP (ln ms<sup>2</sup>) | 7.97 (0.128) | 7.72 (0.187) | 8.2 (0.146) | 7.46 (0.216) | 7.83 (0.133) | 7.06 (0.200) | 7.33 (0.143) | 6.67 (0.248) | 8.08 (0.181) |
| LF/ HF | 1.149 (0.104) | 1.422 (0.150) | 1.411 (0.126) | 1.288 (0.179) | 0.922 (0.110) | 1.386 (0.163) | 0.488 (0.122) | 0.924 (0.210) | 1.203 (0.167) |
| HF<sub>nu</sub> | 0.525 (0.0210) | 0.513 (0.0257) | 0.491 (0.0245) | 0.505 (0.0306) | 0.592 (0.0219) | 0.509 (0.0278) | 0.716 (0.0239) | 0.609 (0.0357) | 0.483 (0.0314) |
| HF<sub>pf</sub> | 0.273 (0.00590) | 0.270 (0.00698) | 0.265 (0.00673) | 0.262 (0.00822) | 0.255 (0.00612) | 0.246 (0.00822) | 0.261 (0.00658) | 0.230 (0.00953) | 0.270 (0.00837) |

*Data are reported as Mean (standard error). Epochs indicated the average number of 3-minute windows being analyzed in each sleep stage.*

**Table 3.2.** Summary of HRV Parameters during QW

| Stage | QW |  |
| --- | --- | --- |
| Age Group (N) | Young (46) | Older (38) |
| Epochs | 14.52 (0.329) | 15.05 (0.362) |
| RR (ms) | 914 (21.3) | 906 (23.4) |
| pNN50 (%) | 31.9 (3.21) | 14.6 (3.53) |
| RMSSD (ln) | 4.09 (0.119) | 3.93 (0.131) |
| HF (ln ms<sup>2</sup>) | 6.43 (0.217) | 6.01 (0.239) |
| LF (ln ms<sup>2</sup>) | 6.72 (0.197) | 6.21 (0.217) |
| TP (ln ms<sup>2</sup>) | 8.19 (0.188) | 7.80 (0.206) |
| LF/ HF | 1.52 (0.192) | 1.92 (0.211) |
| HF<sub>nu</sub> | 0.443 (0.0252) | 0.455 (0.0277) |
| HF<sub>pf</sub> | 0.291 (0.00748) | 0.258 (0.00822) |

*Data are reported as Mean (standard error). Epochs indicated the average number of 3-minute windows being analyzed in each sleep stage.*

### Power spectral analysis

The EEG power spectrum was computed using the Welch method (4 sec Hanning windows with 50 % overlap) (Campbell et al., 2005). Slow-wave activity (SWA; 0.5-1Hz), delta (0.5-4Hz), theta (4-8Hz), alpha (8-13Hz), slow sigma (9-11Hz), and fast sigma (12-15Hz) were calculated for each sleep stage (i.e., WASO, Stage 2, SWS and REM). Power spectrum were averaged across the eight scalp electrodes (F3, F4, C3, C4, P3, P4, O1, O2). Summary statistics for EEG power spectrum are shown in Table 4 for the NAP subjects.

**Table 4.** Summary of EEG Power Across Sleep Stages for the Nap Group

| Stage | WASO |  |  | N2 |  |  | N3 |  |  | REM |  |  |
| --- | --- | --- | --- | --- | --- | --- | --- | --- | --- | --- | --- | --- |
|  | Young | Older |  | Young | Older |  | Young | Older |  | Young | Older |  |
| SWA (0.5-1Hz) | 268.83 (27.70) | 168.19 (39.30) | * | 73.73 (6.10) | 36.94 (3.52) | *** | 169.68 (13.13) | 93.09 (8.98) | *** | 48.07 (16.92) | 26.22 (7.17) | n.s. |
| Delta (0.5-4Hz) | 358.52 (62.34) | 251.41 (36.05) | n.s. | 141.30 (9.37) | 74.80 (5.96) | *** | 308.19 (19.32) | 162.96 (18.45) | *** | 83.86 (25.58) | 48.66 (11.46) | n.s. |
| Theta (4-8Hz) | 32.50 (2.49) | 23.06 (2.85) | * | 10.72 (1.08) | 10.47 (0.68) | n.s. | 14.50 (1.16) | 10.80 (1.68) | n.s. | 10.05 (1.38) | 7.39 (1.57) | n.s. |
| Alpha (8-13 Hz) | 25.29 (3.47) | 29.89 (6.47) | n.s. | 20.03 (0.92) | 15.72 (1.28) | ** | 25.55 (1.43) | 19.08 (2.46) | * | 13.00 (1.83) | 10.37 (2.71) | n.s. |
| Slow Sigma (9-11Hz) | 18.64 (1.51) | 12.10 (1.63) | ** | 5.04 (0.37) | 5.09 (0.56) | n.s. | 5.59 (0.41) | 4.85 (0.72) | n.s. | 4.54 (0.76) | 3.20 (0.71) | n.s. |
| Fast Sigma (12-15Hz) | 10.47 (1.22) | 6.88 (0.91) | * | 4.94 (0.37) | 3.97 (0.30) | * | 5.63 (0.43) | 3.44 (0.44) | *** | 2.32 (0.24) | 2.27 (0.35) | n.s. |

*Data are reported as Mean (standard error).*
*Asterisks indicate significant differences between age groups (n.s. $p > 0.05$ ; \* $p < 0.05$ ; \*\* $p < 0.01$ ; \*\*\* $p < 0.001$ ).*

### Statistical Analyses

All statistical analyses were performed in R 3.6.2, using the libraries lme4 (Bates et al., 2015), and lsmeans (Lenth, 2016).

First, to compare the between-subject difference between pre-nap rest to QW, we used a two-way ANOVA using Stage (pre-nap rest, QW) and Age (Young, Older) as between-subject factors for each HRV variable. We first assessed a reduced (nested) model, with Stage (pre-nap rest, QW) as the only factor. We then included Age (Young, Older) in the full model. By comparing the reduced and full model using the Likelihood Ratio Test (Lewis et al., 2011), we are able to interpret if adding the factor Age significantly improved the model.

Next, in order to investigate within-subject profiles of cardiac activity across sleep stages, we used linear-mixed effect models (LME), which do not depend on limited assumptions about variance-covariance matrix assumptions (sphericity). Additionally, LME models eliminate the need of averaging epochs across sleep stages and allow inclusion of an unbalanced number of observations per subject in the analyses. Moreover, LME models take into account the influence of factors whose levels are extracted randomly from a population (i.e. participants), thus yielding more generalizable results (Baayen et al., 2008). In order to examine the autonomic profile changes across sleep stages in the young adults and older adults separately, we built a model using Participant as crossed random effects and Stage as a fixed effect (Young adult model: prenap-resting, N2, N3, REM, WASO; Older adults model: prenap-resting, N2, N3, WASO) for each HRV variable. Next, to examine the age-related changes in ANS profiles, we built a LME model using Participant as crossed random effects and Stage (prenap-resting, N2, N3, WASO) and Age (Young, Older) as fixed effects. We first built a reduced (nested) model, with Stage (prenap-resting, N2, N3, WASO) as the only fixed effect, and then included Age (Young, Older) as a fixed effect in the full model. Like for the wake analyses, comparing the reduced and full model using the Likelihood Ratio Test (Lewis et al., 2011), we are able to interpret if adding the factor Age significantly improved the model. Tukey’s correction for multiple testing was used for post-hoc comparisons.

To examine the differences in demographics, sleep architecture, and power spectrum between the two age groups, an independent-sample t-test (for continuous variables) and chi-squared independent test (for categorical variables) was performed for each variable and the results are shown in Table 1, Table 2, and Table 4, respectively. Pearson correlation coefficients were used to examine the bivariate relationship between HRV variables of interests and sleep parameters.

## Results

### Demographic, Sleep architecture and Power spectrum Data

As reported in Table 1, there were significant differences between young vs older adults in Age, Ethnicity, caffeine and alcohol intake, ESS, BMI, and habitual bedtime/waketime, while no differences in nap frequency or duration. Older adults reported an earlier bedtime and wake time, higher caffeine and alcohol intake and BMI, as well as lower daytime sleepiness, compared to the young adults.

Furthermore, older adults demonstrated poorer sleep quality during the nap, with significantly less total sleep time and N3 sleep duration. Older adults also presented lower sleep efficiency, more time in N1 sleep, and more wake after sleep onset (WASO) than the younger adults (see Table 2).

In addition, older adults showed lower power in the SWA, Delta, Alpha, and Fast Sigma frequency bands during NREM sleep, compared to the younger adults (see Table 4).

### Comparing Cardiac Profiles during QW and pre-nap rest: Young vs Older Adults

Means and standard errors for the HRV variables during QW are provided in Table 3.2.

We examined the autonomic profile between a QW episode and pre-nap rest for older and younger adults. For RR, we report no significant main effect of Stage (F_(1,185)_ = 2.8112, p = 0.0953), Age (F_(1,185)_ = 0.0114, p = 0.9151), or interaction (F_(1,185)_ = 0.0428, p = 0.8364). The likelihood ratio test was not significant (LR = 0.0271; p = 0.9733), suggesting that adding the factor Age did not significantly improved the model.

For the time-domain measure, RMSSD_ln_ showed no significant Age effect (F_(1,185)_ = 1.2418, p = 0.2666), Stage effect (F_(1,185)_ = 2.1883, p = 0.1408), or interaction effect (F_(1,185)_ = 0.0670, p = 0.7961). The likelihood ratio test was not significant (LR = 0.6544; p = 0.5210), suggesting that adding the factor Age did not significantly improved the model. Again, we did not detect age differences in this variable between QW and pre-nap rest.

For frequency-domain variables, our analysis revealed no significant main effect of Stage (F_(1,185)_ = 0.4507, p = 0.5028), Age (F_(1,185)_ = 2.5930, p = 0.1090), or interaction (F_(1,185)_ = 0.1611, p = 0.6886) for TPln. Similarly, no significant main effect of Stage (F_(1,185)_ = 2.4654, p = 0.1181), Age (F_(1,185)_ = 3.6948, p = 0.0561, or interaction (F_(1,185)_ = 0.0003, p = 0.9857) were found for HFln. Again, for LFln, no significant main effect of Stage (F_(1,185)_ = 0.0135, p = 0.9078), Age (F_(1,185)_ = 3.7253, p = 0.0551, or interaction (F_(1,185)_ = 0.0051, p = 0.9434) were found. None of the likelihood ratio tests were significant (TPln: LR = 1.3771; p = 0.2549; HFln: LR = 1.8476; p = 0.1605; LFln: LR = 1.8652; p = 0.1578), suggesting that adding the factor Age did not improve these models. These results suggested that young and older subjects show similar autonomic profiles in a standard waking HRV condition and a pre-nap rest condition. However, for LF/HF ratio, our analysis revealed a Stage effect (F_(1,185)_ = 5.2704, p = 0.0228), with QW showing a higher ratio than pre-nap resting. No significant Age effect (F_(1,185)_ = 3.0999, p = 0.0800) or interaction effect were found (F_(1,185)_ = 0.1010, p = 0.7510). Similar patterns were shown in HF_nu_, with a Stage effect (F_(1,185)_ = 8.1255, p = 0.0049) where subjects showed higher HF_nu_ during pre-nap resting compared to QW. No significant Age effect (F_(1,185)_ = 0.0027, p = 0.9588) or interaction effect were found (F_(1,185)_ = 0.2267, p = 0.6346). Again, the likelihood ratio test was not significant for either of these dependent measures (LF/HF ration: LR = 1.6005; p = 0.2046; Hfnu: LR = 0.1147; p = 0.8917), suggesting that adding the factor Age did not improved these models. This suggests that the relative proportion of vagally-mediated HF activity is slightly higher directly before a nap compared to a QW period for both older and younger adults.

In summary, by comparing the models, we found that adding the factor of Age did not improve the models of any wake HRV variables, suggesting no evidence of age-related changes in ANS activity during wake using both the standard QW condition and pre-nap rest.

### Cardiac Autonomic Profiles during a Nap in Young Adults

Next, we compared pre-nap rest, sleep stages (N2, N3, REM) and wake after sleep onset (WASO) in younger subjects. Means and standard errors for the HRV variables across sleep stages are provided in Table 3.1.

For RR, we report a significant effect for Stage (F_(4,150)_ = 13.9887, p < 0.0001; Figure 1a), with Tukey’s post-hoc revealing a significant lengthening of RR intervals during N2 and N3 relative to Rest (N2: p = 0.0015; N3: p = 0.0096), WASO (all ps < 0.0001), and REM sleep (all ps < 0.0001), but no differences between N2 and N3 (p = 0.9995), nor between REM and Rest (p = 0.0851) or WASO (p = 0.9397). First, for TP_ln_ we showed a significant Stage effect (F_(4,150)_ = 9.773, p < 0.001; Figure 1e), with Tukey’s post-hoc showing lower power during N3 relative to Rest (p < 0.001), WASO (p < 0.001), REM sleep (p = 0.0006), and N2 sleep (p = 0.0036). This was primarily due to reductions in LF power (F_(4,150)_ = 12.320, p < 0.0001; Figure 1d), with lower LF power emerging during N3 relative to Rest (p < 0.0001), WASO (p < 0.0001), N2 (p < 0.0001), and REM sleep (p < 0.0001), but no shifts in HF_ln_ (F_(4,150)_ = 1.1838, p = 0.3203; Figure 1c) across sleep stages. These patterns were also clear when examining both the LF/HF ratio (F_(4,150)_ = 10.0353, p < 0.0001; Figure 1f) and HFnu (F_(4,150)_ = 23.1608, p < 0.0001; Figure 1g), Tukey’s post-hoc tests revealing significantly lower LF/HF ratio during N3 relative to Rest (p = 0.001), WASO (p < 0.001), REM sleep (p = 0.0021), and N2 sleep (p = 0.0243) and significantly greater HFnu during N3 relative to Rest, WASO, REM, and N2 sleep (all ps < 0.0001). HFnu was also greater during N2 relative to Rest, WASO, and REM (p = 0.002). For the time-domain variable, no differences were found in RMSSD (F_(4,150)_ = 0.968, p = 0.4272; Figure 1b) across stages. Taken together, these results confirm previous studies demonstrating that among younger adults, as the brain shifts from wake to deeper stages of NREM sleep, the body undergoes marked changes with heart rate deceleration and a transition to parasympathetic dominance in ANS activity.

**Figure 1.**
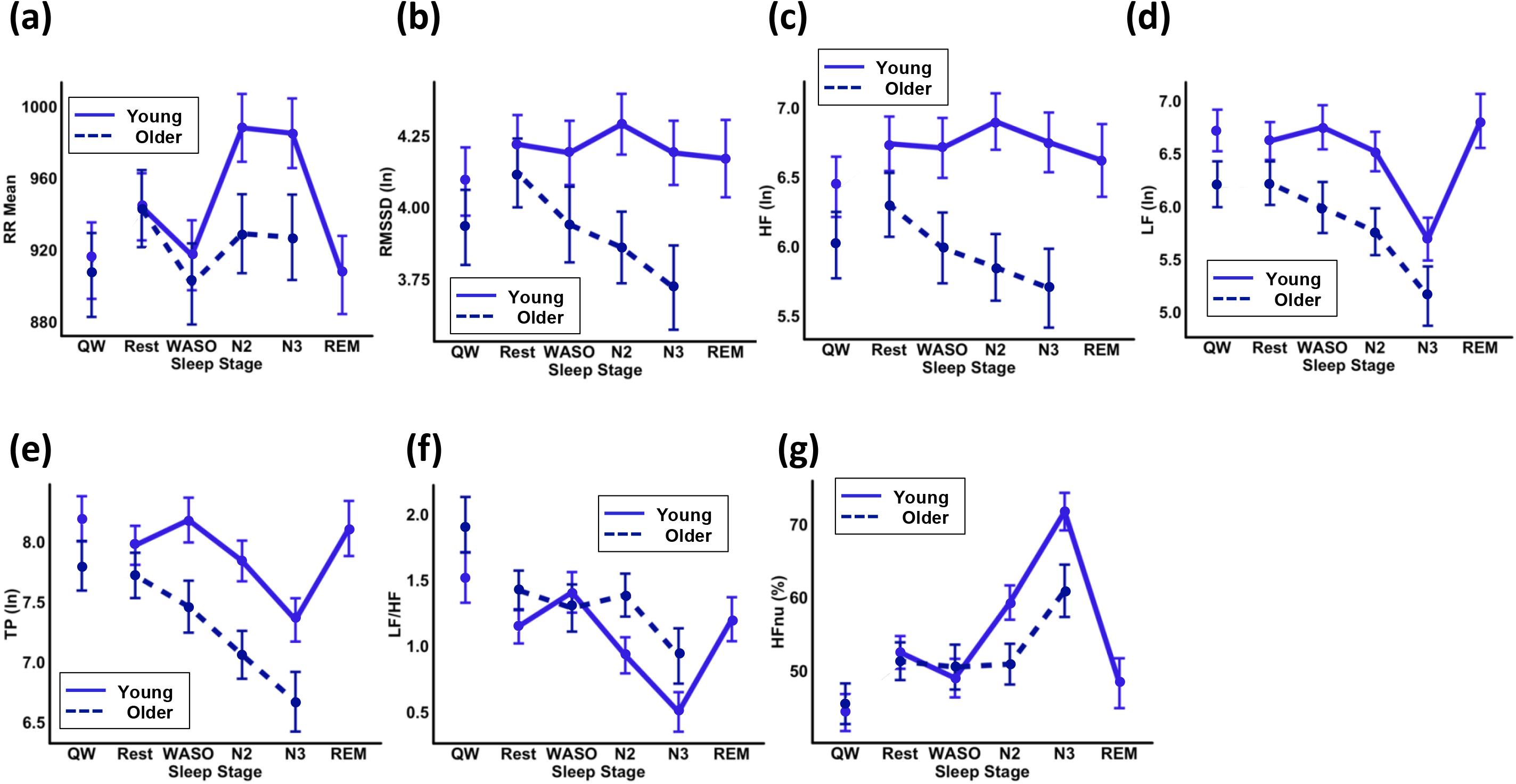
(a–h) Heart rate variability (HRV) components across sleep stages in younger (solid blue line) and older (dashed darkened blue line) adults in the NAP group. (a) Mean of RR intervals (ms); (b) RMSSD (ln); (c) HF HRV (ln); (d) LF HRV (ln); (e) Total Power (f) LF/HF Ratio (g) HFnu. Error bars represent standard error of the mean. Note that the QW and Rest analysis is a between subjects’ comparison.

### Cardiac Autonomic Profiles during a Nap in Older Adults

Next, we compared pre-nap rest, sleep stages (N2, N3, REM) and wake after sleep onset (WASO) in older subjects. For older adults, RR showed a significant effect for Stage (F_(3, 82)_ = 7.5560, p < 0.0001; Figure 1a), with Tukey’s post-hoc showing a significant reduction of RR intervals during WASO relative to Rest (p = 0.0025). But, no differences emerged across the sleep stages. For frequency domain indicators, we showed a significant TP_ln_ Stage effect (F_(3, 82)_ = 4.8486, p = 0.0037; Figure 1e) with significantly lower power during N3 (p = 0.0050) and N2 (0.0364) relative to Rest. Similar to the younger cohort, this was primarily due to reductions in LF power (LF_ln_; F_(3, 82)_ = 2.8704, p = 0.0413; Figure 1d), with significantly lower LF power during N3 relative to Rest (p = 0.0260), but no shifts in HF_ln_ (F_(3, 82)_ = 1.4464, p = 0.2353; Figure 1c) across sleep stages. Unlike the youngers, sleep modulations were not shown in the LF/HF ratio Stage effect (F_(3, 82)_ = 1.3850, p = 0.2532; Figure 1f) or HF_nu_ (F_(3, 82)_ = 2.5453, p = 0.0617; Figure 1g). Additionally, for the time-domain variable, no significant differences were found for RMSSD_ln_ (F_(3, 82)_ = 2.2311, p = 0.0907; Figure 1b) across stages.

Overall, these results implicate a similar pattern in ANS total and LF power across a nap for older and younger adults, whereas we found no evident sleep modulations in HFnu and LF/HF ratio among our older subjects.

### Comparing Cardiac Profiles during Nap: Young vs Older Adults

We utilized linear mixed-model analysis to directly compare profiles across older and younger adults. For RR intervals (Figure 1a), a Stage effect (F_(3,212)_ = 14.901, p < 0.0001), and an Age x Stage interaction emerged (F_(3,212)_ = 5.477, p = 0.0012). Young adults demonstrated a significant lengthening in RR during N2 and N3 relative to Rest (N2: p = 0.0001; N3: p = 0.0015) and WASO (all ps < 0.0001). For older adults, no sleep-dependent shifts were present (all ps>0.1275), however we found reduced RR during WASO compared Rest (p = 0.0062). The likelihood ratio test was significant (LR = 17.3995; p = 0.0016), suggesting that adding the factor Age significantly improved the model.

For frequency-domain variables, we observed main effects of Age for TPln (F_(1,105)_ = 7.215, p = 0.0084; Figure 1e), with the older adults displaying significantly lower total power overall. Furthermore, a main effect of Stage was found for TPln (F_(3,212)_ = 12.302, p < 0.0001; Figure 1e), with significantly lower total power during N3, compared to Rest (p < 0.0001), WASO (p < 0.0001), and N2 (p = 0.0205), as well as significantly lower total power during N2, compared to Rest (p = 0.0083) and WASO (p = 0.0468). No significant main effect of Stage was found (F_(3,212)_ = 1.6810, p = 0.1720). No significant Age x Stage interaction was found (F_(3,212)_ = 1.808, p = 0.1467) for TPln. The likelihood ratio test was significant (LR= 12.5808; p = 0.0135), suggesting that adding the factor Age improved the model.

These shifts in total power were complimented by a significant Age effect for HF_ln_ (F_(1,105)_ = 8.3446, p = 0.0047; Figure 1c) and LF_ln_ (F_(1,105)_ = 7.329, p = 0.0079; Figure 1d), with the older adults showing significantly lower HF and LF power, compared to the younger cohort. In addition, for LF_ln_, a main effect of Stage was found (F_(3,212)_ = 12.283, p < 0.0001), with significantly lower LF power during N3, compared to Rest (p < 0.0001), WASO (p < 0.0001), and N2 (p = 0.0006). No significant main effect of Stage was found for HF_ln_ (F_(3,212)_ = 0.7454, p = 0.5261). No significant Age x Stage interaction was found for HF_ln_ (F_(3,212)_ = 2.3701, p = 0.0716) or LF_ln_ (F_(3,212)_ = 0.654, p = 0.5815). However, the likelihood ratio test was significant for HF_ln_ (LR = 15.2829; p = 0.0041) but not for LF_ln_ (LR = 9.2153; p = 0.0559), suggesting that adding the factor Age significantly improved the explained variation in HF_ln_ but not LF_ln_.

The LF/HF ratio showed a Stage effect (F_(3,212)_ = 9.6908, p < 0.0001; Figure 1f), with significantly lower LF/HF ratio during N3, compared to Rest (p = 0.0002), WASO (p = 0.0003), and N2 (p = 0.0086). No significant Age effect was shown (F_(1,105)_ = 2.9395, p = 0.0894). No significant Age x Stage interaction was found (F_(3,212)_ = 1.7531, p = 0.1572) for LF/HF ratio. In addition, the likelihood ratio test was not significant (LR = 8.2874; p = 0.0816), suggesting that adding the factor Age did not improve the model.

For HF_nu_, a significant effect of Stage (F_(3,212)_ = 23.4128, p < 0.0001; Figure 1g), with significantly greater HF_nu_ during N3, compared to Rest, WASO, and N2 (all ps < 0.0001). No significant Age effect was shown (F_(1,105)_ = 2.3738, p = 0.1264). Here, an Age x Stage interaction was apparent (F_(3,212)_ = 3.0343, p = 0.0302), with young adults expressing significantly greater HFnu during N2 (p = 0.0216) and N3 (p = 0.0158) but not the rest of stages, compared to older adults. The likelihood ratio test was significant (LR = 11.5583; p = 0.021), suggesting that adding the factor Age improved the model.

For the time-domain measure RMSSD, we also observed an Age effect (F_(1,105)_ = 4.330, p = 0.0399; Figure 1b), with older adults showing lower HRV than the youngers overall. Furthermore, an Age x Stage interaction for RMSSD_ln_ were found (F_(3,212)_ = 2.872, p = 0.0373; Figure 1c), with older adults showing significantly lower vagally-mediated HRV during N2 sleep (p = 0.0083) and N3 (p = 0.0084), compared to the younger cohort. The likelihood ratio test was significant (LR = 12.9717; p = 0.0114), suggesting that adding the factor Age significantly improved the model.

In summary, by comparing the LME models, the current study showed that adding the factor of Age improved the models of RR intervals, RMSSD_ln_, HF_ln_, TP_ln_, HF_nu_. In particular, older adults demonstrated overall decreases in TP_ln_, HF_ln_, LF_ln_, compared to the youngers. The Age x Stage interactions were found in HFnu and RMSSD_ln_, with young adults expressing significantly greater vagal activity during NREM sleep, but not during WASO or pre-nap rest, compared to the older cohort. Taken together, these findings suggest an age-related decline with significantly less parasympathetic dominance during a daytime nap in older adults, compared with youngers.

### Relationship between HRV Variables and Sleep parameters

Next, given the parallel declines in parasympathetic activity, sleep architecture, and EEG activity, we further investigate if the age-related differences in HRV variables might be driven by EEG activity or sleep architecture. We examined the correlation between parasympathetic activity (RMSSDln, HFln, HFnu) and sleep parameters. We found no significant correlation between HRV variables and EEG activity (SWA, delta, theta, alpha, slow sigma, and fast sigma) during N2 (Young: all p> 0.1969; Older: all p> 0.6994), N3 (Young: all p> 0.0754; Older: all p> 0.1974), or REM (Young: all p> 0.4447). Furthermore, no significant correlation between HRV variables and sleep architecture: total sleep time (Young: all p> 0.2826; Older: all p> 0.1352), sleep efficiency (Young: all p> 0.4732; Older: all p> 0.3271), N1 sleep duration (Young: all p> 0.1203; Older: all p> 0.3271), N2 sleep duration (Young: all p> 0.1175; Older: all p> 0.3740), N3 sleep duration (Young: all p> 0.1105; Older: all p> 0.0793), WASO (Young: all p> 0.0697; Older: all p> 0.1892). Taken together, these results suggested that the dampened parasympathetic activity during NREM sleep are neither associated nor explained simply by differences in sleep architecture or reductions in EEG activity during the nap.

## Discussion

Heart rate variability reflects an individual’s ability to adapt to moment-to-moment changes in the environment (Thayer & Lane, 2009). In the current study, we aimed to assess the cardiac autonomic activity during a daytime nap in healthy young and older adults. There is compelling evidence for a strong link between decline in cardiac autonomic activity during wake and both cognitive and health outcomes, as well as links between reduced HRV and age-related decreases in physiological functioning. However, the modulation of cardiac ANS activity in this population specifically during a daytime nap, compared with quiet wake, is under-investigated. Here, in agreement with our hypotheses, our young subjects showed parasympathetic dominance during NREM sleep compared to pre-nap resting. In addition, the current study showed that older adults exhibited a reduced autonomic variability, or cardiac autonomic rigidity, during NREM sleep, compared to the younger cohort. Lastly, no age-related modulations were evident during wake, suggesting a sleep-specific reduction in parasympathetic activity in older adults.

### A Nap is a Mini-Cardiovascular Break in Young Adults

In the current work, the cardiac autonomic pattern across daytime sleep in young adults was consistent with the prior literature. Indeed, similar to our results, lengthening of RR intervals has been observed during N2 and N3 compared to Rest, WASO, and REM sleep in studies examining nocturnal sleep (de Zambotti et al., 2014; Trinder et al., 2001), as well as a daytime nap (Cellini et al., 2016) among healthy young adults. Regarding parasympathetic/vagal activity specifically, it’s been previously reported that vagal activity rises at sleep onset and remains elevated across the whole sleep period, with higher vagal modulation during N2 and N3 than REM sleep and waking (Nocturnal Sleep: Ako et al., 2003; de Zambotti et al., 2014; Trinder et al., 2001; Nap: Cellini et al., 2016), suggesting vagal dominance during NREM sleep stages. Similar to previous findings, we found that HF_nu_ was significantly increased during N3 compared to Rest, WASO, N2, and REM, and during N2 compared to Rest, WASO, and REM. These studies suggested that, similar to nighttime sleep, a daytime nap demonstrates dominance of vagally-mediated activity during NREM sleep, in particular during N3.

Indeed, this autonomic shift was also reflected in the LF/HF ratio, which was also at its lowest during N3. A similar pattern was reported, with LF/HF ratio reduced in N3 compared to N2, pre-sleep wakefulness, and REM sleep (Nocturnal Sleep: Ako et al., 2003; Nap: Cellini et al., 2016). We also observed a strong decrease in TP during N3 mainly due to the significant decrease in LF_ln_ power during this sleep stage (HFln was not significantly reduced in N3), similar to previous studies examining nocturnal sleep (de Zambotti et al., 2014) and nap (Cellini et al., 2016). These data suggest discrete fluctuations in LF_ln_ power and an overall dominance of vagal activity during N3 sleep in a daytime nap, which mirror nighttime sleep HRV profiles (Whitehurst et al., 2018). Note that, since the mechanism underlying LF_ln_ is still debated (Billman, 2013; Reyes del Paso et al., 2013), strong inferences about the physiological mechanisms that contribute to changes in LF and LF/HF ratio are difficult to make for these parameters, but we report them for descriptive purposes.

Taken together, these findings are consistent with previous reports on the cardiac autonomic profile during nocturnal sleep and daytime nap in young adults (Ako et al., 2003; Cellini et al., 2016; de Zambotti et al., 2014; Trinder et al., 2001). Collectively, these studies demonstrate an overall reduction in cardiovascular output and dominance of parasympathetic/vagal activity during NREM sleep (Busek et al., 2005), which has been associated with significant benefits for the cardiovascular system (Trinder et al., 2012). Considering that the most common reason for napping in healthy young adults seems to be for restorative purposes (Duggan et al., 2018), these data suggest that daytime naps may provide a compensatory protective window for the cardiovascular system (Naska et al., 2007) and serve as a “mini cardiovascular break” that might potentially benefit psychological and physiological health.

### Reduced Mini-Cardiovascular Break in Older Adults’ Naps

Autonomic imbalance, indexed by low HF HRV and elevated sympathetic activity, is associated with various pathological conditions, such as cardiovascular disease, diabetes, and Alzheimer’s disease. Given the parasympathetic dominance during sleep, nighttime sleep has been described as a “cardiovascular holiday” (Trinder et al., 2012). However, in our older subjects, the expected cardiovascular break was severely dampened, which builds upon previous findings examining nocturnal sleep (Brandenberger et al., 2003; Crasset et al., 2001). We first demonstrated no age differences between two waking conditions: a standard measure of quiet wake in which subjects lie in bed not intending to sleep, and a pre-nap rest condition more typical of sleep studies. These data are important for comparing across studies but also show that older adults show the same parasympathetic profile during wake as younger adults.

Our examination of autonomic profiles across pre-nap rest, wake after sleep onset and sleep stages revealed that older adults may not confer the same restorative benefits from a nap as younger adults. We reported a drop in absolute HRV indices during NREM sleep specifically, in older subjects, compared with young subjects, with shorter RR intervals, lower TPln, lower vagally-mediated HRV (reflected by RMSSDln, HFln, and HFnu). The decreased vagally-mediated HRV in aging reveals a tendency for a relative increase in vagal withdrawal and a predominant loss of parasympathetic activity, which may be related to the increased number of awakenings during the nap and lower duration of N3 in this age group (see Table 1). Whether the impaired autonomic balance causes the N3 decrease, or whether the decrease in N3 is simply reflected in vagally-mediated HRV, is not yet known. Overall, the current findings are in alignment with previous studies comparing younger vs older adult HRV during nocturnal sleep (Brandenberger et al., 2003). Although the total power dropped drastically from Rest to N2 and N3 in our elderly subjects, the relative increase in parasympathetic activity during N3 (reflected by HF_nu_) suggested that the elderly might still have cardiovascular benefits from napping; however, nap might not as restorative as their younger counterparts. Noteworthy, the age-related decrease in vagally-mediated HRV were only shown in the nap but not in either wake conditions, suggesting a differential loss in ANS activity that is sleep-specific among older subjects compared to the youngers. Taken together, HRV during naps, but not quiet wake, might be a novel biomarker or target of intervention for cardiovascular health.

Although there is now substantial evidence that low HRV values during wake are associated with increased mortality risk in the elderly (Tsuji et al. 1994), it is not known if a drop HRV indices during NREM sleep would be an additional risk factor for adverse cardiac events. Future studies on sleep interventions are needed to better elucidate the relationship between the dynamic, bidirectional association between sleep and the autonomic system. Intriguingly, a recent study showed that acoustic enhancement of SWA increased parasympathetic activity (measured by HFnu) in healthy young adults during nocturnal sleep (Grimaldi et al., 2019). Given that our data provide further evidence of parasympathetic loss during NREM sleep in aging, a variety of sleep intervention techniques (i.e. acoustic stimulation, non-invasive brain stimulation (NIBS) techniques) could potentially be utilized to consolidate N3 and enhance cardiovascular benefits or cognition in older populations. Furthermore, a number of studies have shown that physical training increases HRV in older subjects (Raffin et al., 2019). It can then be suggested that regular exercise might benefit the autonomic imbalance in the elderly and would restore the association between HRV and NREM sleep. Taken together, these findings have implications for translational medicine, and suggest that aging can represent a potentially informative model to investigate the complex interaction between autonomic activity, sleep, and health.

### Limitations

The present study is not without limitations. First, our sample was a convenience sample of both men and women, and uncontrolled variations in hormonal status among young women can have an impact on cardiac vagal activity (see Schmalenberger et al., 2019 for a review). Although time-domain measures (i.e., RMSSD) tend to be more stable across menstrual cycle, frequency-domain measures of HRV may be susceptible to hormonal fluctuation (Sato et al., 1995; Yazar & Yazıcı, 2016). Future studies controlling for hormonal fluctuation will be needed to confirm the age-related changes in ANS profiles during naps. As a second limitation, no standardized diagnostic assessment of psychiatric disorder or chronic medical disease was performed to confirm the self-reported screening surveys. Thirdly, older subjects showed higher BMI, higher caffeine and alcohol intake than the younger cohorts. Although none of our participants reported a clinically significant medical condition or substance dependence, a subtle impact of obesity, caffeine, and alcohol on HRV cannot be excluded. Furthermore, older subjects expressed fragmented sleep architecture compared to the youngers, with lower sleep efficiency, greater wake after sleep onset, less duration of SWS, and lower power in the slow oscillation, delta, and fast sigma bands during NREM sleep. However, we found no significant correlation between HRV variables and sleep parameters during any sleep stages. Additionally, we cannot rule out that there are individual differences in circadian cycling and the midday nap times that we chose for both older and younger adults likely aligned with different phases in individual’s natural circadian cycling. This may have influenced our results as the circadian system is known to impact on HRV. In addition, we also analyzed the frequency peak of HF (HFfp) in order to control for respiratory rate, which can affect the HRV (Song & Lehrer, 2003). HFfp showed no difference between the two age groups and varied within a narrow range in the HF spectrum, between 0.23 and 0.29 Hz, with the greatest difference of about 3 breaths per minute. Thus, the respiratory activity does not seem to play a key role in the age-related differences of HRV modulation. However, we cannot completely exclude that our younger and older subjects have no alteration in the cardiopulmonary coupling, as may be detected using measures of coherence (Thomas et al., 2005). Lastly, standard practice for HRV analyses (Task Force, 1996) requires assessment of HRV over a 5-minute period of a consistent sleep stage (longer than 3-min in the current study). However, due to the large amount of sleep transitions present in a daytime nap, especially in older adults, this method decreased the number of available subjects and may have biased the sample towards less fragmented sleepers. Notably, several studies have presented data with shorter epoch windows and demonstrated robust results (Shaffer & Ginsberg, 2017; Whitehurst et al., 2018). Nevertheless, we cannot exclude the impact of shorter windows on measures of HRV modulation.

### Conclusions

The current study is the first, to our knowledge, to compare ANS profiles between young versus older adults during a daytime nap and a period of quiet resting. This is clinically relevant, given that nap frequency increases with age and that changes in HRV are associated with cardiovascular mortality. In summary, an increase in vagal withdrawal and a predominant loss of parasympathetic activity during NREM sleep in a daytime nap were found in our older subjects, compared to the youngers, while not such age-related changes in autonomic profiles were found during quiet wake.

## Acknowledgements

This work was supported by the National Institutes of Health R01 (AG046646) and (AG061355). P.C. analyzed data and wrote the paper; S.C.M, L.N.W. and N.S. designed the study; L.N.W. and N.S. performed research; and S.C.M. supervised research. S.C.M, L.N.W. and N.S. edited the paper.

## References

Ako, M., Kawara, T., Uchida, S., Miyazaki, S., Nishihara, K., Mukai, J., Hirao, K., Ako, J., & Okubo, Y. (2003). Correlation between electroencephalography and heart rate variability during sleep. Psychiatry and Clinical Neurosciences, 57(1), 59–65. https://doi.org/10.1046/j.1440-1819.2003.01080.x

Baayen, R. H., Davidson, D. J., & Bates, D. M. (2008). Mixed-effects modeling with crossed random effects for subjects and items. Special Issue: Emerging Data Analysis, 59(4), 390–412. https://doi.org/10.1016/j.jml.2007.12.005

Bates, D., Mächler, M., Bolker, B., & Walker, S. (2015). Fitting Linear Mixed-Effects Models Using lme4. Journal of Statistical Software, Articles, 67(1), 1–48. https://doi.org/10.18637/jss.v067.i01

Benjamin, E. J., Blaha, M. J., Chiuve, S. E., Cushman, M., Das, S. R., Deo, R., Floyd, J., Fornage, M., Gillespie, C., Isasi, C., & others. (2017). Heart disease and stroke statistics-2017 update: A report from the American Heart Association. Circulation, 135(10), e146–e603.

Billman, G. E. (2013). The LF/HF ratio does not accurately measure cardiac sympatho-vagal balance. Frontiers in Physiology, 4, 26–26. PubMed. https://doi.org/10.3389/fphys.2013.00026

Blackwell, T., Yaffe, K., Ancoli-Israel, S., Schneider, J. L., Cauley, J. A., Hillier, T. A., Fink, H. A., Stone, K. L., & for the Study of Osteoporotic Fractures Group. (2006). Poor Sleep Is Associated With Impaired Cognitive Function in Older Women: The Study of Osteoporotic Fractures. The Journals of Gerontology: Series A, 61(4), 405–410. https://doi.org/10.1093/gerona/61.4.405

Boardman, A., Schlindwein, F. S., Rocha, A. P., & Leite, A. (2002). A study on the optimum order of autoregressive models for heart rate variability. Physiological Measurement, 23(2), 325–336. https://doi.org/10.1088/0967-3334/23/2/308

Borb, A. A., & Achermann, P. (1999). Sleep Homeostasis and Models of Sleep Regulation. Journal of Biological Rhythms, 14(6), 559–570. https://doi.org/10.1177/074873099129000894

Brandenberger, G., Ehrhart, J., Piquard, F., & Simon, C. (2001). Inverse coupling between ultradian oscillations in delta wave activity and heart rate variability during sleep. Clinical Neurophysiology, 112(6), 992–996. https://doi.org/10.1016/S1388-2457(01)00507-7

Brandenberger, G., Viola, A. U., Ehrhart, J., Charloux, A., Geny, B., Piquard, F., & Simon, C. (2003). Age-related changes in cardiac autonomic control during sleep. Journal of Sleep Research, 12(3), 173–180. https://doi.org/10.1046/j.1365-2869.2003.00353.x

Burr, R. L. (2007). Interpretation of normalized spectral heart rate variability indices in sleep research: A critical review. Sleep, 30(7), 913–919. PubMed.

Campbell, S. S., Murphy, P. J., & Stauble, T. N. (2005). Effects of a Nap on Nighttime Sleep and Waking Function in Older Subjects. Journal of the American Geriatrics Society, 53(1), 48–53. https://doi.org/10.1111/j.1532-5415.2005.53009.x

Cellini, N., Whitehurst, L. N., McDevitt, E. A., & Mednick, S. C. (2016). Heart rate variability during daytime naps in healthy adults: Autonomic profile and short-term reliability. Psychophysiology, 53(4), 473–481. PubMed. https://doi.org/10.1111/psyp.12595

Chadda, K. R., Ajijola, O. A., Vaseghi, M., Shivkumar, K., Huang, C. L.-H., & Jeevaratnam, K. (2018). Ageing, the autonomic nervous system and arrhythmia: From brain to heart. Ageing Research Reviews, 48, 40–50. https://doi.org/10.1016/j.arr.2018.09.005

Chen, P.-C., Whitehurst, L. N., Naji, M., & Mednick, S. C. (2020). Autonomic Activity during a Daytime Nap Facilitates Working Memory Improvement. Journal of Cognitive Neuroscience, 1–12. https://doi.org/10.1162/jocn_a_01588

Chung, F., F. R. C. P. C., Yegneswaran, B., M. B. B. S., Liao, P., M. D., Chung, S. A., Ph. D., Vairavanathan, S., M. B. B. S., Islam, S., M. Sc., Khajehdehi, A., M. D., & Shapiro, C. M., F. R. C. P. C. (2008). STOP Questionnaire: A Tool to Screen Patients for Obstructive Sleep Apnea. Anesthesiology: The Journal of the American Society of Anesthesiologists, 108(5), 812–821. https://doi.org/10.1097/ALN.0b013e31816d83e4

Colosimo, A., Giuliani, A., Mancini, A. M., Piccirillo, G., & Marigliano, V. (1997). Estimating a cardiac age by means of heart rate variability. American Journal of Physiology-Heart and Circulatory Physiology, 273(4), H1841–H1847. https://doi.org/10.1152/ajpheart.1997.273.4.H1841

Crasset, V., Mezzetti Silvia, Antoine Martine, Linkowski Paul, Degaute Jean Paul, & van de Borne Philippe. (2001). Effects of Aging and Cardiac Denervation on Heart Rate Variability During Sleep. Circulation, 103(1), 84–88. https://doi.org/10.1161/01.CIR.103.1.84

Cross, N., Terpening, Z., Rogers, N. L., Duffy, S. L., Hickie, I. B., Lewis, S. J. G., & Naismith, S. (2015). Napping in older people ‘at risk’ of dementia: Relationships with depression, cognition, medical burden and sleep quality. Journal of Sleep Research, 24(5), 494–502. https://doi.org/10.1111/jsr.12313

de Zambotti, M., Cellini, N., Baker, F. C., Colrain, I. M., Sarlo, M., & Stegagno, L. (2014). Nocturnal cardiac autonomic profile in young primary insomniacs and good sleepers. International Journal of Psychophysiology, 93(3), 332–339. https://doi.org/10.1016/j.ijpsycho.2014.06.014

Duggan, K. A., McDevitt, E. A., Whitehurst, L. N., & Mednick, S. C. (2018). To Nap, Perchance to DREAM: A Factor Analysis of College Students’ Self-Reported Reasons for Napping. Behavioral Sleep Medicine, 16(2), 135–153. PubMed. https://doi.org/10.1080/15402002.2016.1178115

Electrophysiology Task Force of the European Society of Cardiology the North American Society of Pacing. (1996). Heart Rate Variability. Circulation, 93(5), 1043–1065. https://doi.org/10.1161/01.CIR.93.5.1043

Gatz, M., Reynolds, C. A., John, R., Johansson, B., Mortimer, J. A., & Pedersen, N. L. (2002). Telephone screening to identify potential dementia cases in a population-based sample of older adults. International Psychogeriatrics, 14(3), 273–289. https://doi.org/10.1017/S1041610202008475

Grimaldi, D., Papalambros, N. A., Reid, K. J., Abbott, S. M., Malkani, R. G., Gendy, M., Iwanaszko, M., Braun, R. I., Sanchez, D. J., Paller, K. A., & Zee, P. C. (2019). Strengthening sleep–autonomic interaction via acoustic enhancement of slow oscillations. Sleep, 42(zsz036). https://doi.org/10.1093/sleep/zsz036

Guo, Y.-F., & Stein, P. K. (2003). Circadian rhythm in the cardiovascular system: Chronocardiology. American Heart Journal, 145(5), 779–786.

Hansen, A. L., Johnsen, B. H., Sollers, J. J., Stenvik, K., & Thayer, J. F. (2004). Heart rate variability and its relation to prefrontal cognitive function: The effects of training and detraining. European Journal of Applied Physiology, 93(3), 263–272. https://doi.org/10.1007/s00421-004-1208-0

Hansen, A. L., Johnsen, B. H., & Thayer, J. F. (2003). Vagal influence on working memory and attention. International Journal of Psychophysiology, 48(3), 263–274. https://doi.org/10.1016/S0167-8760(03)00073-4

Heckenberg, R. A., Eddy, P., Kent, S., & Wright, B. J. (2018). Do workplace-based mindfulness meditation programs improve physiological indices of stress? A systematic review and meta-analysis. Journal of Psychosomatic Research, 114, 62–71. https://doi.org/10.1016/j.jpsychores.2018.09.010

Horne, J. A., & Östberg, O. (1976). A self-assessment questionnaire to determine morningness-eveningness in human circadian rhythms. International Journal of Chronobiology.

Jarczok, M. N., Guendel, H., McGrath, J. J., & Balint, E. M. (2019). Circadian Rhythms of the Autonomic Nervous System: Scientific Implication and Practical Implementation. In Chronobiology-The Science of Biological Time Structure. IntechOpen.

Jasper. (1958). The Ten-Twenty Electrode System of the International Federation. Electroencephalography and Clinical Neurophysiology, 10(371–375).

Johns, M. W. (1991). A new method for measuring daytime sleepiness: The Epworth Sleepiness Scale. Sleep: Journal of Sleep Research & Sleep Medicine, 14(6), 540–545. https://doi.org/10.1093/sleep/14.6.540

Keage, H. A. D., Banks, S., Yang, K. L., Morgan, K., Brayne, C., & Matthews, F. E. (2012). What sleep characteristics predict cognitive decline in the elderly? Sleep Medicine, 13(7), 886–892. https://doi.org/10.1016/j.sleep.2012.02.003

Kemp, A. H., & Quintana, D. S. (2013). The relationship between mental and physical health: Insights from the study of heart rate variability. Psychophysiology in Australasia-ASP Conference-November 28-30 2012, 89(3), 288–296. https://doi.org/10.1016/j.ijpsycho.2013.06.018

Krygier, J. R., Heathers, J. A. J., Shahrestani, S., Abbott, M., Gross, J. J., & Kemp, A. H. (2013). Mindfulness meditation, well-being, and heart rate variability: A preliminary investigation into the impact of intensive Vipassana meditation. Psychophysiology in Australasia-ASP Conference-November 28-30 2012, 89(3), 305–313. https://doi.org/10.1016/j.ijpsycho.2013.06.017

Laborde, S., Mosley, E., & Thayer, J. F. (2017). Heart Rate Variability and Cardiac Vagal Tone in Psychophysiological Research – Recommendations for Experiment Planning, Data Analysis, and Data Reporting. Frontiers in Psychology, 8, 213. https://doi.org/10.3389/fpsyg.2017.00213

Lenth, R. (2016). Least-Squares Means: The R Package lsmeans. Journal of Statistical Software, Articles, 69(1), 1–33. https://doi.org/10.18637/jss.v069.i01

Lewis, F., Butler, A., & Gilbert, L. (2011). A unified approach to model selection using the likelihood ratio test. Methods in Ecology and Evolution, 2(2), 155–162. https://doi.org/10.1111/j.2041-210X.2010.00063.x

Lin, L.-C., Chiang, C.-T., Lee, M.-W., Mok, H.-K., Yang, Y.-H., Wu, H.-C., Tsai, C.-L., & Yang, R.-C. (2013). Parasympathetic activation is involved in reducing epileptiform discharges when listening to Mozart music. Clinical Neurophysiology, 124(8), 1528–1535. https://doi.org/10.1016/j.clinph.2013.02.021

Mankus, A. M., Aldao, A., Kerns, C., Mayville, E. W., & Mennin, D. S. (2013). Mindfulness and heart rate variability in individuals with high and low generalized anxiety symptoms. Behaviour Research and Therapy, 51(7), 386–391. https://doi.org/10.1016/j.brat.2013.03.005

Mattarozzi, K., Colonnello, V., Thayer, J. F., & Ottaviani, C. (2019). Trusting your heart: Long-term memory for bad and good people is influenced by resting vagal tone. Consciousness and Cognition, 75, 102810. https://doi.org/10.1016/j.concog.2019.102810

McEwen, B. S. (2002). Sex, stress and the hippocampus: Allostasis, allostatic load and the aging process. Brain Aging: Identifying the Brakes and Accelerators, 23(5), 921–939. https://doi.org/10.1016/S0197-4580(02)00027-1

Mednick, S. C., McDevitt, E. A., Walsh, J. K., Wamsley, E., Paulus, M., Kanady, J. C., & Drummond, S. P. A. (2013). The Critical Role of Sleep Spindles in Hippocampal-Dependent Memory: A Pharmacology Study. The Journal of Neuroscience, 33(10), 4494. https://doi.org/10.1523/JNEUROSCI.3127-12.2013

Mednick, S., Nakayama, K., & Stickgold, R. (2003). Sleep-dependent learning: A nap is as good as a night. Nature Neuroscience, 6(7), 697–698. https://doi.org/10.1038/nn1078

Monk, T. H. (2005). Aging Human Circadian Rhythms: Conventional Wisdom May Not Always Be Right. Journal of Biological Rhythms, 20(4), 366–374. https://doi.org/10.1177/0748730405277378

Mosley, E., Laborde, S., & Kavanagh, E. (2018). Coping related variables, cardiac vagal activity and working memory performance under pressure. Acta Psychologica, 191, 179–189. https://doi.org/10.1016/j.actpsy.2018.09.007

Naska, A., Oikonomou, E., Trichopoulou, A., Psaltopoulou, T., & Trichopoulos, D. (2007). Siesta in Healthy Adults and Coronary Mortality in the General Population. Archives of Internal Medicine, 167(3), 296–301. https://doi.org/10.1001/archinte.167.3.296

O’Brien, I. A., O’Hare, P., & Corrall, R. J. (1986). Heart rate variability in healthy subjects: Effect of age and the derivation of normal ranges for tests of autonomic function. British Heart Journal, 55(4), 348–354. PubMed. https://doi.org/10.1136/hrt.55.4.348

Ohayon, M. M., & Vecchierini, M.-F. (2002). Daytime Sleepiness and Cognitive Impairment in the Elderly Population. Archives of Internal Medicine, 162(2), 201–208. https://doi.org/10.1001/archinte.162.2.201

Ohayon, M. M., & Zulley, J. (1999). Prevalence of naps in the general population. Sleep and Hypnosis, 1(2), 88–97.

Pavlov, V. A., & Tracey, K. J. (2012). The vagus nerve and the inflammatory reflex—Linking immunity and metabolism. Nature Reviews Endocrinology, 8(12), 743–754. https://doi.org/10.1038/nrendo.2012.189

Pinna, G. D., Maestri, R., Torunski, A., Danilowicz-Szymanowicz, L., Szwoch, M., La Rovere, L. T., & Raczak, G. (2007). Heart rate variability measures: A fresh look at reliability. Clinical Science, 113(3), 131–140. https://doi.org/10.1042/CS20070055

Raffin, J., Barthélémy, J.-C., Dupré, C., Pichot, V., Berger, M., Féasson, L., Busso, T., Da Costa, A., Colvez, A., Montuy-Coquard, C., Bouvier, R., Bongue, B., Roche, F., & Hupin, D. (2019). Exercise Frequency Determines Heart Rate Variability Gains in Older People: A Meta-Analysis and Meta-Regression. Sports Medicine, 49(5), 719–729. https://doi.org/10.1007/s40279-019-01097-7

Reyes del Paso, G. A., Langewitz, W., Mulder, L. J. M., van Roon, A., & Duschek, S. (2013). The utility of low frequency heart rate variability as an index of sympathetic cardiac tone: A review with emphasis on a reanalysis of previous studies. Psychophysiology, 50(5), 477–487. https://doi.org/10.1111/psyp.12027

Russoniello, C. (2013). Heart Rate Variability and Biological Age: Implications for Health and Gaming. Cyberpsychology, Behavior, and Social Networking, 16, 302–308. https://doi.org/10.1089/cyber.2013.1505

Santillo, E., Migale, M., Fallavollita, L., Marini, L., & Balestrini, F. (2012). Electrocardiographic Analysis of Heart Rate Variability in Aging Heart. https://doi.org/10.5772/22538

Sato, N., Miyake, S., Akatsu, J., & Kumashiro, M. (1995). Power Spectral Analysis of Heart Rate Variability in Healthy Young Women During the Normal Menstrual Cycle. Psychosomatic Medicine, 57(4). https://journals.lww.com/psychosomaticmedicine/Fulltext/1995/07000/Power_Spectral_Analysis_of_Heart_Rate_Variability.4.aspx

Schmalenberger, K. M., Eisenlohr-Moul, T. A., Würth, L., Schneider, E., Thayer, J. F., Ditzen, B., & Jarczok, M. N. (2019). A Systematic Review and Meta-Analysis of Within-Person Changes in Cardiac Vagal Activity across the Menstrual Cycle: Implications for Female Health and Future Studies. Journal of Clinical Medicine, 8(11), 1946. PubMed. https://doi.org/10.3390/jcm8111946

Shaffer, F., & Ginsberg, J. P. (2017). An Overview of Heart Rate Variability Metrics and Norms. Frontiers in Public Health, 5, 258–258. PubMed. https://doi.org/10.3389/fpubh.2017.00258

Sinnreich, R., Kark, J. D., Friedlander, Y., Sapoznikov, D., & Luria, M. H. (1998). Five minute recordings of heart rate variability for population studies: Repeatability and age–sex characteristics. Heart, 80(2), 156. https://doi.org/10.1136/hrt.80.2.156

Smith, R., Thayer, J. F., Khalsa, S. S., & Lane, R. D. (2017). The hierarchical basis of neurovisceral integration. Neuroscience & Biobehavioral Reviews, 75, 274–296. https://doi.org/10.1016/j.neubiorev.2017.02.003

Song, H.-S., & Lehrer, P. M. (2003). The effects of specific respiratory rates on heart rate and heart rate variability. Applied Psychophysiology and Biofeedback, 28(1), 13–23.

Thayer, J. F., & Lane, R. D. (2007). The role of vagal function in the risk for cardiovascular disease and mortality. Special Issue of Biological Psychology on Cardiac Vagal Control, Emotion, Psychopathology, and Health., 74(2), 224–242. https://doi.org/10.1016/j.biopsycho.2005.11.013

Thayer, J. F., & Lane, R. D. (2009). Claude Bernard and the heart–brain connection: Further elaboration of a model of neurovisceral integration. The Inevitable Link between Heart and Behavior: New Insights from Biomedical Research and Implications for Clinical Practice, 33(2), 81–88. https://doi.org/10.1016/j.neubiorev.2008.08.004

Thayer, J. F., Yamamoto, S. S., & Brosschot, J. F. (2010). The relationship of autonomic imbalance, heart rate variability and cardiovascular disease risk factors. International Journal of Cardiology, 141(2), 122–131. https://doi.org/10.1016/j.ijcard.2009.09.543

Thomas, R. J., Mietus, J. E., Peng, C.-K., & Goldberger, A. L. (2005). An electrocardiogram-based technique to assess cardiopulmonary coupling during sleep. Sleep, 28(9), 1151–1161.

Tobaldini, E., Nobili, L., Strada, S., Casali, K., Braghiroli, A., & Montano, N. (2013). Heart rate variability in normal and pathological sleep. Frontiers in Physiology, 4, 294. https://doi.org/10.3389/fphys.2013.00294

Trinder, J., Kleiman, J., Carrington, M., Smith, S., Breen, S., Tan, N., & Kim, Y. (2001). Autonomic activity during human sleep as a function of time and sleep stage. Journal of Sleep Research, 10(4), 253–264. https://doi.org/10.1046/j.1365-2869.2001.00263.x

Trinder, J., Waloszek, J., Woods, M. J., & Jordan, A. S. (2012). Sleep and cardiovascular regulation. Pflügers Archiv-European Journal of Physiology, 463(1), 161–168. https://doi.org/10.1007/s00424-011-1041-3

Whitehurst, L. N., Cellini, N., McDevitt, E. A., Duggan, K. A., & Mednick, S. C. (2016). Autonomic activity during sleep predicts memory consolidation in humans. Proceedings of the National Academy of Sciences, 113(26), 7272. https://doi.org/10.1073/pnas.1518202113

Whitehurst, L. N., Chen, P.-C., Naji, M., & Mednick, S. C. (2020). New directions in sleep and memory research: The role of autonomic activity. Current Opinion in Behavioral Sciences, 33, 17–24. https://doi.org/10.1016/j.cobeha.2019.11.001

Whitehurst, L. N., Naji, M., & Mednick, S. C. (2018). Comparing the cardiac autonomic activity profile of daytime naps and nighttime sleep. Neurobiology of Sleep and Circadian Rhythms, 5, 52–57. https://doi.org/10.1016/j.nbscr.2018.03.001

Williams, D. P., Cash, C., Rankin, C., Bernardi, A., Koenig, J., & Thayer, J. F. (2015). Resting heart rate variability predicts self-reported difficulties in emotion regulation: A focus on different facets of emotion regulation. Frontiers in Psychology, 6, 261. https://doi.org/10.3389/fpsyg.2015.00261

Yazar, Ş., & Yazıcı, M. (2016). Impact of Menstrual Cycle on Cardiac Autonomic Function Assessed by Heart Rate Variability and Heart Rate Recovery. Medical Principles and Practice ◻: International Journal of the Kuwait University, Health Science Centre, 25(4), 374–377. PubMed. https://doi.org/10.1159/000444322

Zhang, J. (2007). Effect of Age and Sex on Heart Rate Variability in Healthy Subjects. Journal of Manipulative and Physiological Therapeutics, 30(5), 374–379. https://doi.org/10.1016/j.jmpt.2007.04.001

